# nextPARS: Parallel probing of RNA structures in Illumina

**DOI:** 10.1101/174144

**Authors:** Ester Saus, Jesse Willis, Leszek P. Pryszcz, Heinz Himmelbauer, Toni Gabaldón

## Abstract

RNA molecules play important roles in virtually every cellular process. These functions are often mediated through the adoption of specific structures that enable RNAs to interact with other molecules. Thus, determining the secondary structures of RNAs is central to understanding their function and evolution. In recent years several sequencing-based approaches have been developed that allow probing structural features of thousands of RNA molecules present in a sample. Here, we describe nextPARS, a novel Illumina-based implementation of *in-vitro* parallel probing of RNA structures. Our approach achieves comparable accuracy to previous implementations, while enabling higher throughput and sample multiplexing.

## INTRODUCTION

The structure of RNA molecules plays a central role in their function and regulation (Cruz and Westhof 2009). Over the last years several new approaches have been developed that couple high-throughput sequencing technologies to traditional enzymatic- or chemically-based assays to probe RNA structure, thereby enabling the structural profiling of transcribed RNAs at genome-wide scales (Table 1). These include, among others, PARS (Kertesz et al. 2010), FragSeq (Underwood et al. 2010) or the more recent *in vivo* approaches DMS-Seq (Rouskin et al. 2014; Ding et al. 2014), icSHAPE (Spitale et al. 2015) and PARIS (Lu et al. 2016). Analysis *in vivo* represents a powerful tool to validate *in vitro* studies and to obtain more physiologically relevant information. However, currently available *in vivo* methods can only probe either single-stranded or double-stranded regions, but not both at the same time unless using different technologies, missing the direct combined information obtained in PARS, for example. In addition, RNA is generally associated with proteins and other molecules, which limits the obtained information to unprotected regions (Ge and Zhang 2015). Thus, both *in vitro* and *in vivo* studies are complementary and there is a niche of applications for the two approaches (Sanbonmatsu 2016). In this context, efforts have been made to develop computational methods that infer RNA secondary structure by accounting for such high-throughput experimental data (Ouyang et al. 2013; Wu et al. 2015; Ge and Zhang 2015).

**Table 1.** Summary and main characteristics of methods to probe RNA secondary structure. Main methods available for probing RNA secondary structure. Columns indicate, in this order: the name of the method; the sequencing platform; the possibility of multiplexing different samples in the same lane when sequencing; whether the method tests RNA *in vivo* or *in vitro*; whether single-stranded, double-stranded or both types of RNA regions are probed; the possibility of running the experiments at genome-wide scales; whether the methodology is based on enzymatic or chemical probing. Abbreviations: ds, double-stranded; ss, single-stranded.

| Method | Sequencing platform | Multiplex | *In vitro* - *In vivo* | Secondary structure detected | Genome-wide | Probing | Reference |
| --- | --- | --- | --- | --- | --- | --- | --- |
| **nextPARS** | Illumina | yes | *in vitro* | ds / ss | yes | enzymatic |  |
| **PARS** | Solid | yes | *in vitro* | ds / ss | yes | enzymatic | (Kertesz et al. 2010) |
| **PARS adapted** | Illumina | no | *in vitro* | ds / ss | yes | enzymatic | (Wan et al. 2013) |
| **FragSeq** | Solid | yes | *in vitro* | ss | yes | enzymatic | (Underwood et al. 2010) |
| **SHAPE-Seq** | Illumina | yes | *in vitro* | ss | no | chemical | (Lucks et al. 2011) |
| **SAHPES** | Illumina | yes | *in vitro* | ss | no | chemical | (Poulsen et al. 2015) |
| **MAP-Seq** | Illumina | yes | *in vitro* | ss | no | chemical | (Seetin et al. 2014) |
| **DMS-Seq** | Illumina | yes | *in vivo* | A/C in ss | yes | chemical | (Rouskin et al. 2014; Ding et al. 2014) |
| **icSHAPE** | Illumina | yes | *in vivo* | ss | yes | chemical | (Spitale et al. 2015) |
| **PARIS** | Illumina | yes | *in vivo* | ds | yes | chemical | (Lu et al. 2016) |

Abbreviations: ds, double-stranded; ss, single-stranded.

Here, we describe nextPARS, an adaptation of the PARS technique - originally developed using SOLiD sequencing - to Illumina technology, which allows higher throughput and sample multiplexing. Although the PARS approach has been previously adapted to Illumina (Wan et al. 2013), that protocol does not enable pooling of different samples and, moreover, it requires the use of an Ambion kit which has been discontinued (https://www.lifetechnologies.com/order/catalog/product/4454073). As a consequence, the use of that protocol has been limited to very few studies (Dominissini et al. 2016; Wan et al. 2014; Righetti et al. 2016). Here we developed a related method, based on the parallel specific enzymatic digestion of single-stranded (S1) and double-stranded (V1) regions directly followed by Illumina library preparation and sequencing. Among all previously published methods for probing RNA secondary structure *in vitro*, nextPARS is the only one capable of tagging all four bases in a genome-wide scale, while enabling multiplexing in Illumina’s sequencing technology, therefore dramatically reducing the sequencing costs and enabling higher throughput. We tested the validity of our results by comparing them with reported RNA structures obtained using similar techniques. We probed poly-adenylated (polyA) and total RNA of *Saccharomyces cerevisiae* as well as various *in vitro*-transcribed RNAs added in the experiments. To estimate the accuracy of our approach we compared our high-throughput results with structural data obtained by crystallography of five RNA molecules, totaling 5,606 bases, including the *Tetrahymena* ribozyme fragment TETp4p6, and the *Saccharomyces cerevisiae* ribosomal RNAs RDN5-1, RDN18-1, RDN25-1 and RDN58-1.

## RESULTS

With the aim to have higher sequencing throughput and multiplexing capacity to study the secondary structure of RNA molecules at a genome-wide scale, we implemented and adapted the PARS protocol (Kertesz et al. 2010) to the Illumina sequencing platform. We refer to this approach as “nextPARS” (see Materials and Methods, and Figure 1 for a more detailed description of the protocol). In brief, our adapted protocol couples initial phosphatase and kinase treatments after RNAse probing step, to allow ligation of the corresponding 5’ and 3’ adaptors of the *Illumina TruSeq Small RNA Sample Preparation Kit.* Then a reverse transcription of the RNA fragments and a PCR amplification are performed to obtain sequencing-ready libraries. Finally, single end read sequencing of the gel size-selected libraries and subsequent mapping allows determining the enzymatic cleavage points at base resolution.

**Figure 1.**
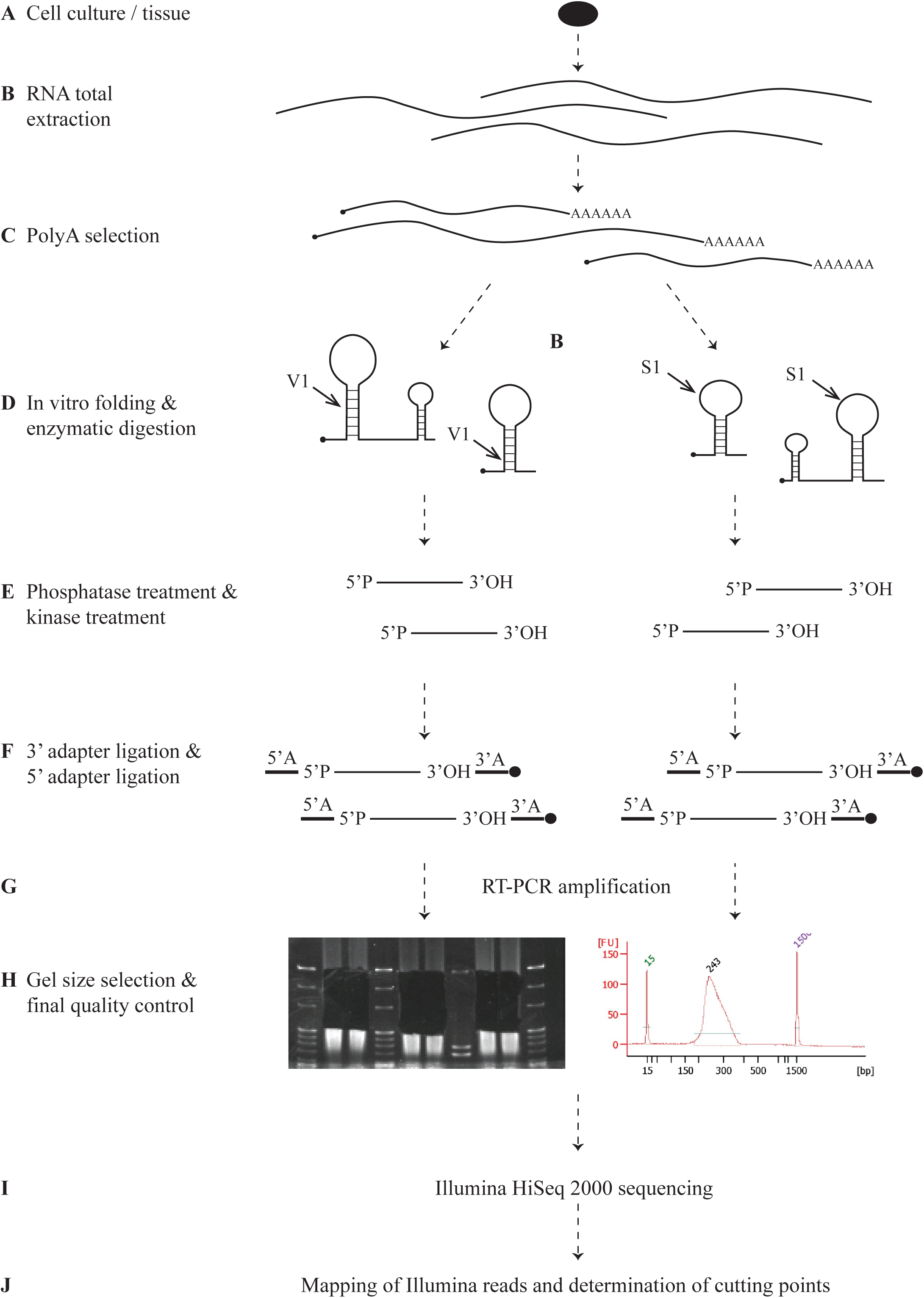
Summary of the different steps performed in the nextPARS protocol. From the cells or tissue of interest **(A)**, total RNA is extracted **(B)** and then polyA+ RNA is selected **(C)** to initially prepare the samples for nextPARS analyses. Once the quality and quantity of polyA+ RNA samples is confirmed, RNA samples are denatured and *in vitro* folded to perform the enzymatic probing of the molecules with the corresponding concentrations of RNase V1 and S1 nuclease **(D)**. For the library preparation using *Illumina TruSeq Small RNA Sample Preparation Kit,* an initial phosphatase treatment of the 3’ends and a kinase treatment of the 5’ ends are required **(E)** to then ligate the corresponding 5’ and 3’ adapters at the ends of the RNA fragments **(F)**. Then a reverse transcription of the RNA fragments and a PCR amplification are performed to obtain the library **(G)**. The library is size-selected to get rid of primers and adapters dimers using an acrylamide gel and a final quality control is performed **(H)**. Libraries are sequenced in single-reads with read lengths of 50 nucleotides using Illumina sequencing platforms **(I)** and computational analyses are done as described in Materials and Methods section in order to map Illumina reads and determine the enzymatic cleavage points **(J)**.

We obtained highly reproducible results in terms of enzymatic digestion profiles, with high and significant statistical correlation among at least three independent replicas (Table 2, Supplementary table S1). These correlations were of the same level as those found in the original PARS protocol. Detailed analysis of the digestion profiles in the control molecules showed significant agreement with published structures, classical chemical footprinting, and results from other methodologies, particularly the SOLiD-based, PARS method (Figure 2). However, we noted that different probing approaches, differences in the relative abundance of molecules, and even the use of different sequencing protocols (e.g. Illumina vs SOLiD) had an impact on reactivity profiles. As a result, results from PARS and nextPARS were positively correlated but showed moderate levels of agreement. Thus raw data provided by both methods should not be considered equivalent but rather related.

**Table 2.** Correlations within nextPARS replicates, within PARS replicates, and between nextPARS and PARS. Correlation values here are the average Spearman’s correlation of the read counts at each site of the corresponding transcripts, comparing all experimental runs in the indicated groups one to one (values in upper triangles of Supplementary Tables S1A and S1B). To be considered, a transcript must have a minimum load of 5.0 (an average read count per site of at least 5.0) in both experimental runs being compared. Average number of comparisons refers to the average number of transcripts between two experimental runs being compared that are found and pass the load threshold (lower triangles of S1A and S1B). Controls refer to the three molecules used as positive controls in the enzymatic probing (TETp4p6, TETp9p9.1, and HOT2).

|  | All yeast + controls |  | Controls |  |
| --- | --- | --- | --- | --- |
|  | Correlation<br>(p-val) | Avg. # of<br>comparisons | Correlation<br>(p-val) | Avg. # of<br>comparisons |
| Within nextPARS, S1 | 0.845 (0.0) | 287.67 | 0.940 (0.0) | 2.33 |
| Within nextPARS, V1 | 0.853 (0.0) | 331.27 | 0.925 (0.0) | 3.00 |
| Within PARS, S1 | 0.857 (0.0) | 219.78 | 0.863 (0.0) | 3.00 |
| Within PARS, V1 | 0.844 (0.0) | 162.93 | 0.753 (0.0) | 3.00 |
| Between nextPARS and PARS, S1 | 0.424 (0.0) | 199.04 | 0.254 (0.0) | 3.00 |
| Between nextPARS and PARS, V1 | 0.459 (0.0) | 170.58 | 0.192 (0.0) | 3.00 |

**Figure 2.**
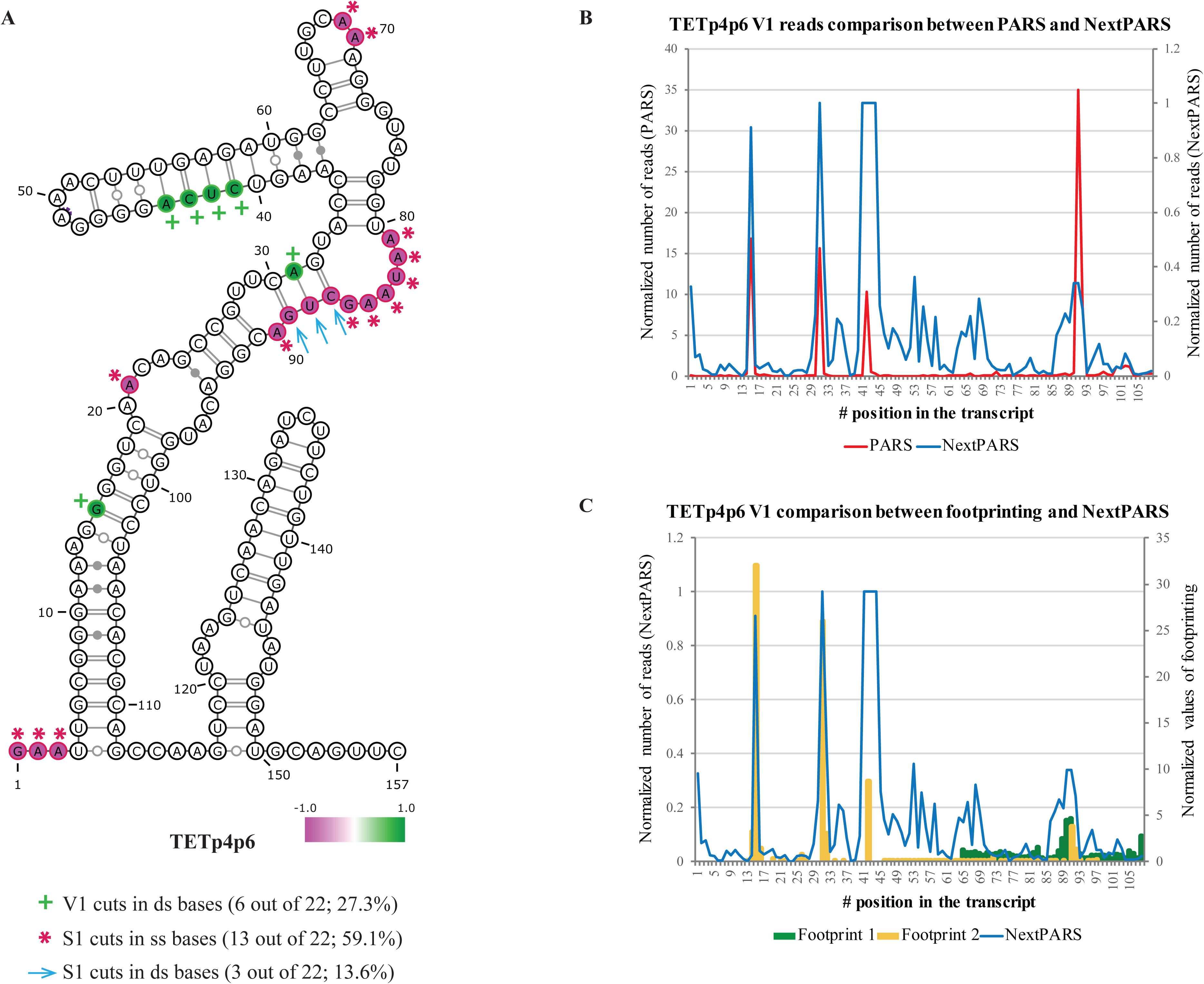
nextPARS results in TETp4p6 fragment. **(A)** Sites having a nextPars score higher that 0.8 (predicted paired site) or lower than -0.8 (predicted unpaired site) are indicated as green and purple, respectively, on the reference secondary structure of TETp4p6 RNA according to PDB database and visualized using VARNA program (Visualization Applet for RNA, http://var-na.lri.fr/, (Darty et al. 2009)). Green crosses (**+**) show V1 cuts (paired sites) which target doublestranded nucleotides in the reference structure, pink asterisks (*) show S1 cuts (unpaired sites) which target single-stranded nucleotides in the reference structure, while blue arrows (**→**) show S1 cuts wrongly targeting double-stranded bases according to the reference structure. For three bases out of 22 (13.6%) we obtained an incorrect signal according to PDB available structure.Noticeably, these bases are located between two loops, which could account for coexistence of different RNA secondary structures with small differences between them for the same molecule.**(B)** Plot comparing both PARS and nextPARS techniques: normalized number of reads for V1 enzyme are plotted for each technique. **(C)** Plot comparing the results obtained with nextPARS with those of previously published results obtained by traditional footprinting experiments (Kertesz et al. 2010).

To provide a better framework for comparison of both methods, we compared PARS and nextPARS results with structural data obtained by crystallography of five RNA molecules, totaling 5,356 bases, including the *Tetrahymena* ribozyme fragment TETp4p6, and the yeast ribosomal RNAs RDN5-1, RDN18-1, RDN25-1 and RDN58-1. The structural data was taken from the Protein Data Bank (PDB), which has files containing 3D coordinates of the molecules within a structure (Berman et al. 2000). TETp4p6 can be found with the PDB ID 1GID, and its structure was determined by Cate, et al. (Cate et al. 1996). The four ribosomal RNAs can be found with the PDB ID 4V88, which is a collection of multiple older PDB IDs -- 3U5H contains RDN5-1, RDN25-1, RDN58-1 as distinct chains, and 3U5F is RDN18-1. The structure of these four rRNAs was determined by Ben-Shem, et al. (Ben-Shem et al. 2011). The original method for calculating the PARS scores was based on the log2 ratio between normalized values of V1 reads and S1 reads for a given PARS experiment (Kertesz et al. 2010). However, due to discrepancies in the read counts between V1 and S1 experiments, as mentioned in the methods sections under “Computation of nextPARS scores” part (iv), we found that this method was unreliable and inconsistent (Supplementary figure S1). We thus used an alternative procedure, which we tested using the set of control molecules mentioned above. The resulting procedure (see Materials and Methods) process the raw digestions profiles so that a single nextPARS score, ranging from -1.0 (highest preference for single-stranded regions) to 1.0 (highest preference for double-stranded regions) is produced. The full computational pipeline to derive these scores from raw nextPARS data can be found at (https://github.com/Gabaldonlab/nextPARS).

We compared results obtained using both scoring methods (PARS and nextPARS) on both sequencing datasets (those provided in the original PARS publication, and those produced here). Our results indicate that both methods are comparable in their sensitivities and accuracy to directly determine single or double-stranded sites, regardless of the scoring scheme used (Figure 3). In fact, when comparing the nextPARS sequencing data and scoring method to those of PARS, we see statistically significant correlations (Spearman coefficient = 0.563). As expected, the correlations are stronger when comparing the two scoring methods on the same sequencing dataset, since the experimental and sequencing methods differ. Correlating the two different scoring methods on the nextPARS dataset gives a coefficient of 0.835, and on the PARS dataset gives a coefficient of 0.874.

**Figure 3.**
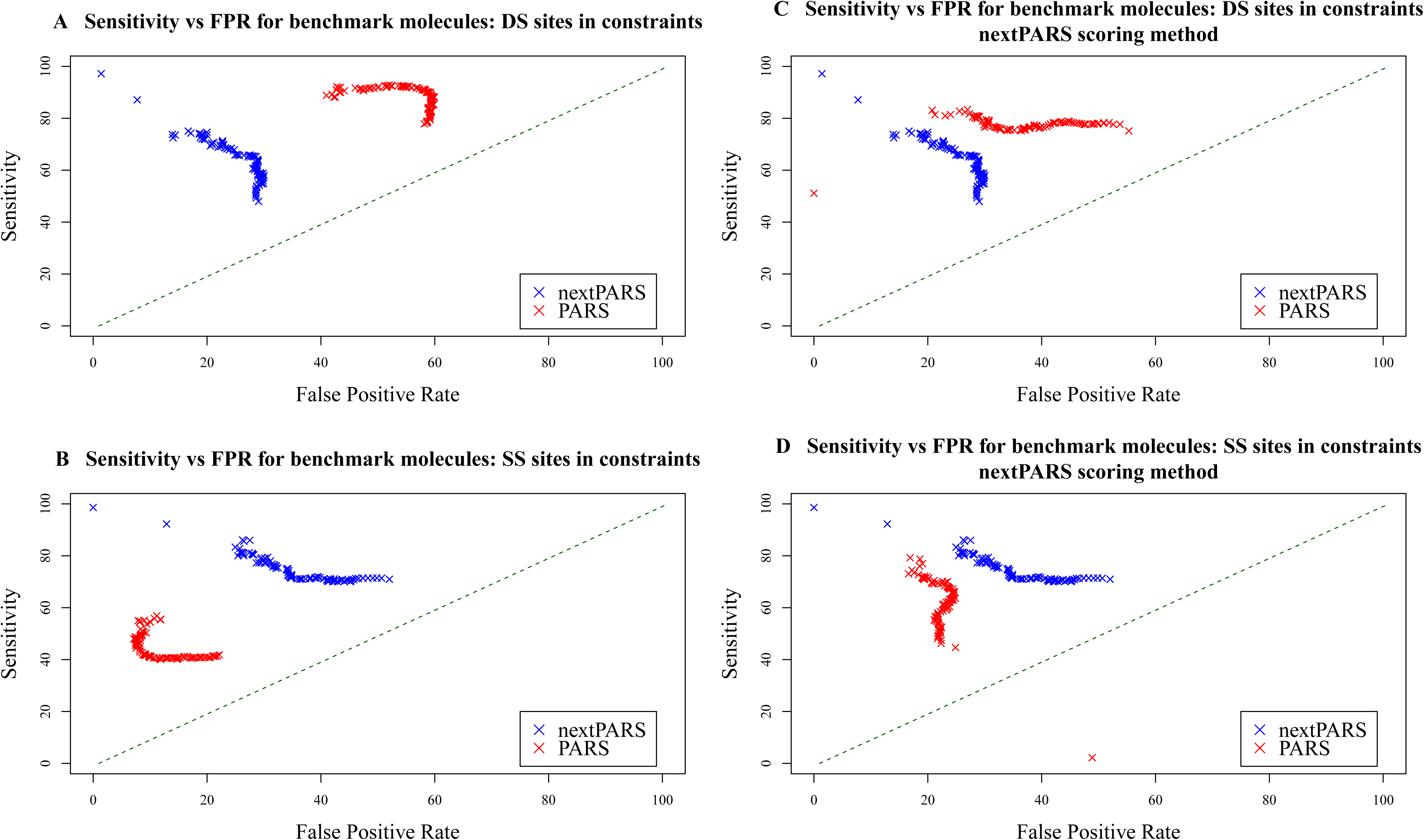
Comparison of scoring methods from nextPARS and PARS. Each panel shows the sensitivity and false positive rate (FPR) of the structural constraints (paired or unpaired) predicted for the indicated data set and scoring method, where each point is the value obtained with a given score threshold. For each scoring method, a range of 100 thresholds is applied to the absolute value of the score in order to test the effect of constraining sites with scores of varying magnitudes; for nextPARS scores, the range is from 0 to 1, which is the maximum score; for PARS scores, since they do not have a fixed range, it is from 0 to the maximum absolute value of the scores. Sensitivity and FPR are determined by comparing constraints to the reference structures for those sites with scores above the given threshold. The three potential constraint values are paired (site has a score greater than or equal to threshold), unpaired (site has a negative score less than or equal to threshold * -1.0), or NA (below threshold, not considered). All values are the average of the 5 control structure molecules, weighted by molecule length. **(A)** Sensitivity and FPR for paired (DS: doublestranded) sites according to the reference. nextPARS data (blue) was scored using the nextPARS scoring method, and PARS data (red) using the PARS scoring method. **(B)** Unpaired (SS: single-stranded) sites according to the reference. Again, nextPARS data was scored using the nextPARS scoring method, and PARS data using the PARS scoring method. **(C)** Paired sites according to the reference. Here, both nextPARS and PARS data were scored using the nextPARS scoring methods to verify that its effect is consistent. **(D)** Unpaired sites according to the references. Again, both nextPARS and PARS data scored using the nextPARS methods.

However, despite this similarity, the nextPARS scoring approach seems to provide more consistent results. When comparing Figure 3A to 3B, 3C to 3D, and Supplementary Figure S1A to S1B, we can see that the ratio of sensitivity to false positive rate (FPR) is more stable between paired and unpaired sites when employing nextPARS scoring than PARS scoring. The red points in 3A and S1A (PARS scoring of paried sites for PARS data and nextPARS, respectively) differ greatly from the red points in 3B and S1B (PARS scoring for unpaired sites), while the respective blue points (nextPARS scoring) remain relatively consistent. Figures 3C and 3D (both red and blue points using nextPARS scoring) show greater similarity between the paired and unpaired sites. This indicates that nextPARS scoring avoids the bias that may be present due to differences in the raw data between the V1 and S1 experiments. In addition, when comparing the accuracy of structures produced (whether a site is correctly predicted to be paired or unpaired) using PARS and nextPARS scoring as constraints (with a score threshold of +/- 0.8), nextPARS had significantly higher accuracy overall (p-value = 0.005) (Figure 4).

**Figure 4.**
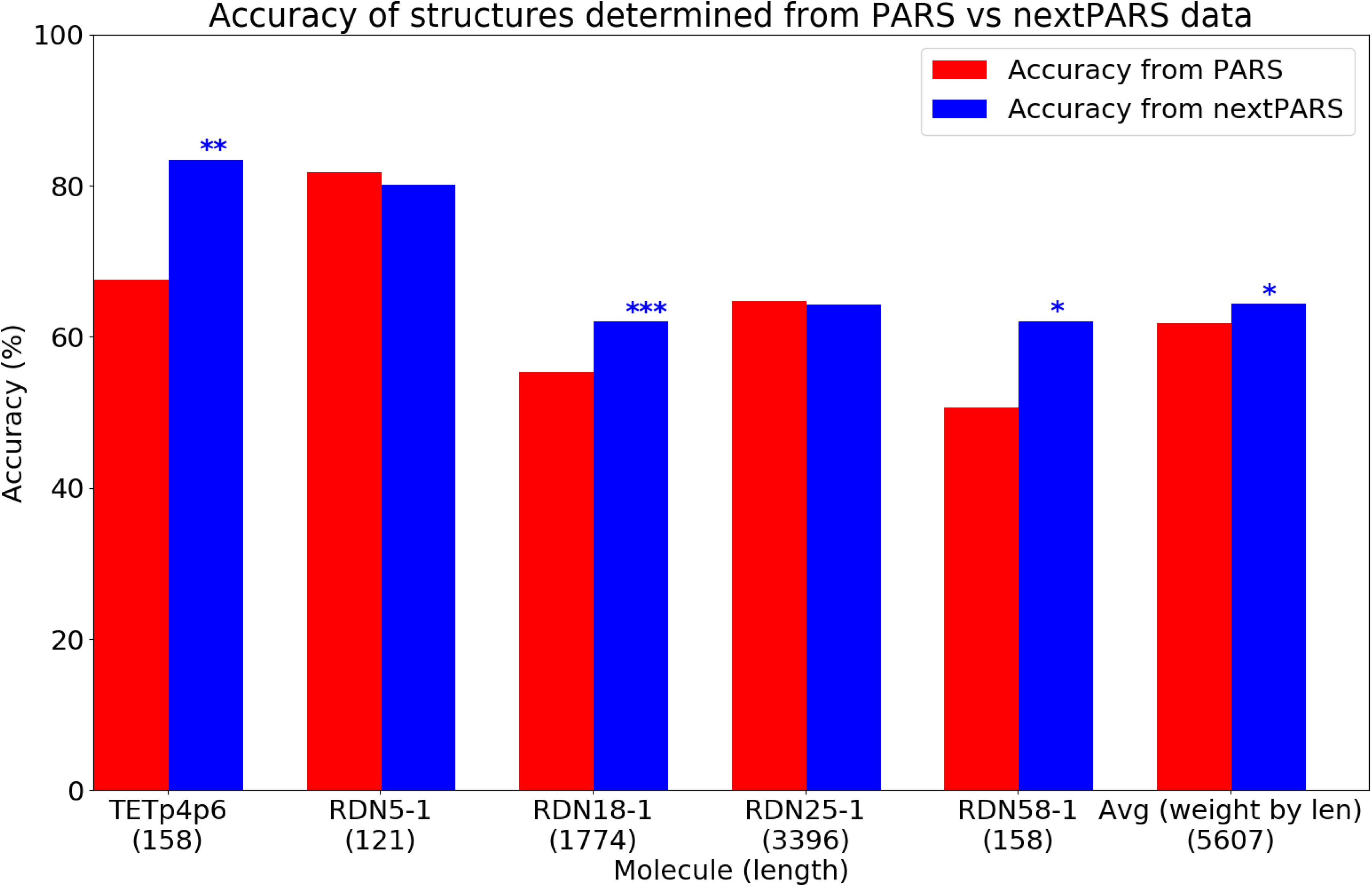
Accuracy of constrained RNAfold structures (the percentage of positions in the molecule which were correctly determined to be either double-stranded or single-stranded). Red bars are values for outputs with PARS scores as constraints, blue bars have nextPARS scores as constraints. The final pair of bars is the average accuracy of the 5 molecules as weighted by the length of each molecule. The stars indicate the level of significant increase of one method over the other (*: p<0.05, **: p<0.005, ***: p<0.0005).

Finally, to test whether stochastic, non-enzymatic breaks could be a source of noise in our protocol, we probed a set of five RNA molecules with known secondary structure with our nextPARS protocol but using RNase A, an enzyme that specifically cuts in single-stranded Cs and Us. We obtained only one cut in an unexpected base (A) out of 79 (Figure 5). Then, differences in signal strengths could account for real cutting preferences of the enzymes, differences in accessibility, or the presence of alternative co-existing structures.

**Figure 5.**
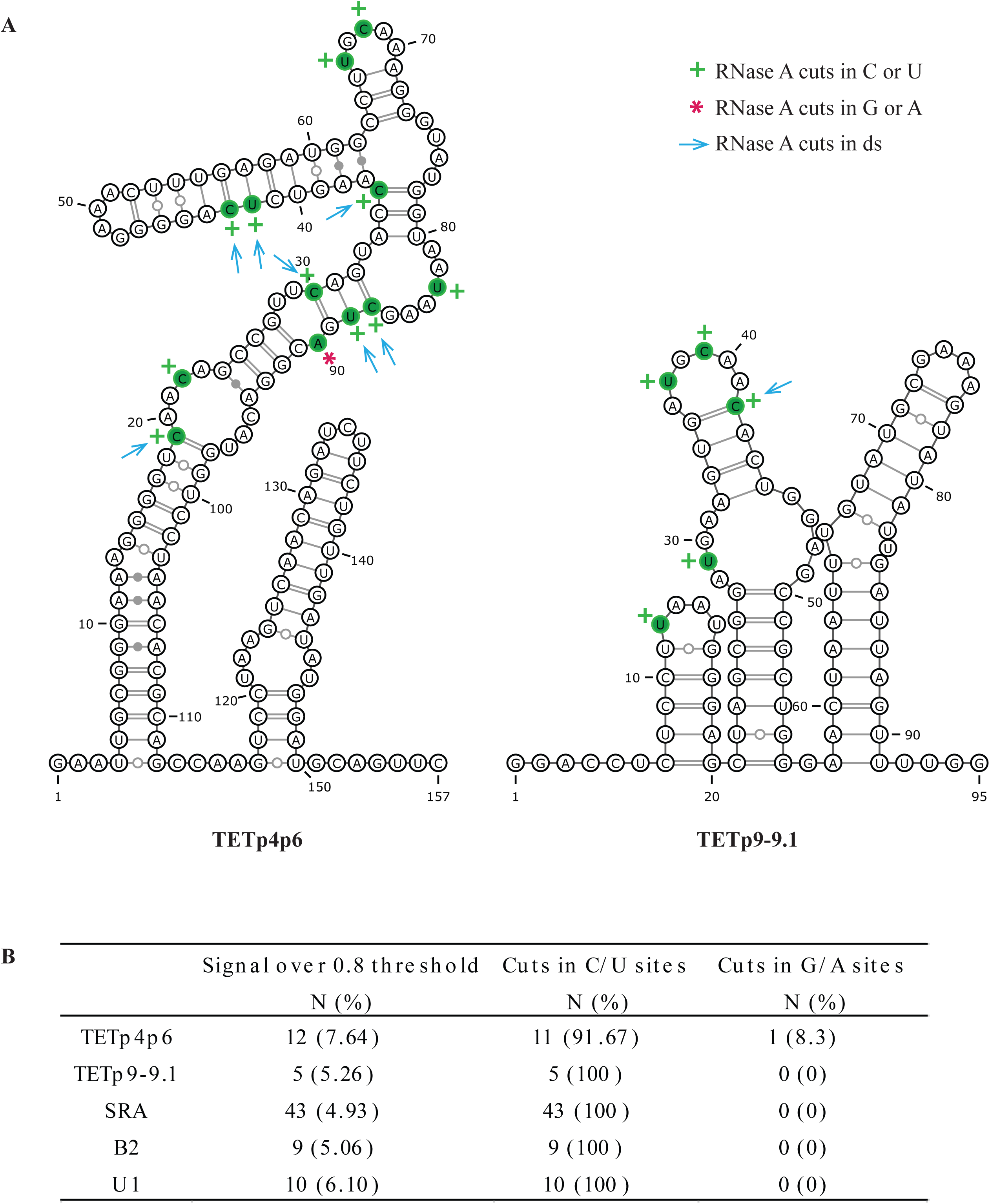
Probing of RNA molecules with RNase A enzyme. Examples of the signals obtained in some RNA molecules when performing nextPARS using RNase A, an enzyme that cuts specifically in single-stranded cytosines (C) and uracils (U). Scores were calculated for each site by first capping all read counts for a given transcript at the 95th percentile and then normalizing to have a maximum of 1 (as done in the “Computation of nextPARS scores” of the Materials and Methods, but since Rnase A is the only enzyme in this case, there will be no subtraction performed, so all values will then fall in the range of 0 to 1). Cuts are considered for signals above a threshold of 0.8. **(A)** nextPARS signals above the threshold of 0.8 are depicted for TETp4p6 and TETp9-9.1 RNA fragments after probing them by nextPARS using RNase A. Secondary structures of the RNA fragments according to PDB are displayed using VARNA program (Visualization Applet for RNA, http://varna.lri.fr/, (Darty et al. 2009)). In green, nucleotides with a cut signal above 0.8; green crosses (+) show cuts obtained in a C or U; pink asterisks (*) show cuts obtained in a G or A; and blue arrows (→) show cuts obtained in double-stranded positions. **(B)** Table summarizing the total number (N) and percentages (%) of cuts witha signal above 0.8 threshold obtained in five different RNA fragments with known secondary structure (TETp4p6, TETp9-9.1, SRA, B2, U1): first column, N and % of cuts with a signal above 0.8 in the molecules; second column, N and % of these cuts in C or U nucleotides; and third column, N and % of cuts in G or A nucleotides.

## DISCUSSION

In this work, we present nextPARS, an adapted method of PARS technology to Illumina platform for RNA structure probing. The main advantages of nextPARS are that the experimental procedure is very straightforward to follow, all scripts are freely available to easily obtain the final secondary structure scores for posterior analyses, and the results are at least as accurate as previous methodologies but allowing higher throughput and sample multiplexing in comparison to PARS.

One of the main limitations in the experimental protocol of nextPARS is that when using the Illumina TruSeq Small RNA kit to prepare the libraries we cannot directly ligate first the 5’adaptor and then the 3’adaptor, requiring the performance of phosphatase and kinase treatments of the RNA ends. This implies that we cannot discern 5’ ends in the RNA molecules caused by V1 or S1 enzymatic digestion from those produced by unspecific fragmentation of RNA molecules (RNase V1 and S1 nuclease enzymatically digest RNA leaving 5’ phosphate ends, while random fragmentation produces 5’-hydroxyl groups which cannot be ligated). In the original PARS protocol (Kertesz et al. 2010; Wan et al. 2013), after the initial enzymatic digestion, a random fragmentation step is done followed by the 5’adaptor ligation that only occurs in 5’-phospate ends. In nextPARS protocol, we skipped the initial random fragmentation after the V1 and S1 specific enzymatic digestion (we obtained high quality libraries and a final gel size-selection is performed) and proceed directly to the adaptors ligations (previous required phosphatase and kinase treatments are done). Thus, it is possible that non-specific fragmentation occurring in RNA molecules during nextPARS protocol would produce noisy signals. However, when enzymatically digesting with RNase A (which specifically cuts in single-stranded Cs and Us) some control molecules with previously described secondary structure, the signals obtained are all in the expected nucleotides except 1 (in one A) out of 79 (Figure 5). This means we can rule out unspecific fragmentation as a possible cause of noisy signals, which could account for the presence of different conformations of RNA molecules at the same time or different enzymatic preferences for cutting points. Besides, we cannot discard that more than one cut occurs in some RNA molecules, which could also lead to noisy signals, since possible conformational changes occurring in the RNA molecule due to the first cut could be detected by a second cut. Although this is an intrinsic characteristic of both PARS and nextPARS techniques that cannot be avoided, we tried to minimize this confounding effect. First, we performed several nextPARS experiments testing different enzyme concentrations, and we chose the optimal V1 and S1 amounts to have single-cut kinetics based on the results obtained for some molecules with known secondary structure. Moreover, data analyses and thresholds are applied to remove as much as possible the noisy signals, to have in the end the more accurate and reliable results possible.

Altogether, nextPARS is a rapid and easy protocol using Illumina sequencing technology to experimentally and massively probe the secondary structure of RNAs. It achieves the same level of resolution, as well as similar accuracy of previously published *in-vitro* structure probing methodologies, while providing higher throughput and multiplexing capacity. In addition, we provide a computational procedure to go from the sequencing reads to a single score that can be used in downstream analyses.

## MATERIALS AND METHODS

### Sample preparation

#### Total and PolyA+ RNA from yeast

*Saccharomyces cerevisiae* S288C was grown in YPDA medium in an orbital shaker (30°C, 200 rpm, overnight). Total RNA from these cultures was extracted using the RiboPure™-Yeast Kit according to manufacturer’s instructions (Ambion), starting with a total amount of 3x10^8^ cells per sample as recommended for a maximum yield. Total RNA integrity and quantity of the samples were assessed using the Agilent 2100 bioanalyzer with the RNA 6000 Nano LabChip® Kit (Agilent, see Supplementary figure S2A) and NanoDrop 1000 Spectrophotometer (Thermo Scientific). To obtain PolyA+ RNA, total RNA from yeast was purified by two rounds of selection using MicroPoly(A)Purist™ Kit according to manufacturer’s instructions (Ambion) to obtain poly(A) RNA from yeast, and the quality of the samples was controlled as above (Supplementary figure S2A).

#### RNA positive controls

Three RNA fragments with previously determined secondary structures were spiked into the samples and used as positive controls in the experiments. *Tetrahymena* ribozyme and HOTAIR clones were obtained from Howard Chang’s lab (Stanford University School of Medicine). Both of them had been previously used as positive controls in the original PARS protocol (Kertesz et al. 2010) and were used in the present work to compare PARS with nextPARS performance. In addition, three other RNA molecules with previously described structures (SRA, B2 and U1) were spiked into the samples in one experiment to probe them with RNase A. In all cases, plasmids were transformed in One Shot® TOP10 Chemically Competent *E. coli* according to manufacturer’s instructions (Invitrogen™). Single colonies were grown in LB+Ampicillin medium (37°C, 220 rpm, overnight), and plasmids were purified using QIAprep Spin Miniprep kit according to the manufacturer’s instructions (Qiagen). PCRs were performed to amplify two fragments of *Tetrahymena* ribozyme (TETp4p6 and TETp9-9.1) and one fragment of the other molecules (HOTAIR -HOT2-, SRA, B2 and U1). Primer sequences and amplicon sizes are shown in Supplementary table S2. PCR amplicons were purified using a QIAquick PCR purification kit according to manufacturer’s instructions (Qiagen) and then were sequenced with Sanger, to confirm that no mutation had been introduced in the fragments of interest. Then, the fragments used as positive controls were transcribed *in vitro* using the T7 RiboMax Large-scale RNA production system according to manufacturer’s instructions (Promega). Finally, RNAs of interest were selected by size and purified using Novex-TBE Urea gels according to manufacturer’s instructions (Life Technologies). A final quality control of the purified RNAs was performed as described above.

### Enzymatic probing

For the enzymatic probing of RNA samples, we reproduced the PARS protocol using RNase V1 and S1 nuclease to cleave RNAs in double or single-stranded conformation, respectively (Kertesz et al. 2010; Wan et al. 2013). 2 μg of PolyA+ RNA or total RNA were mixed with 20 ng of each positive control RNA (TETp4p6, TETp9-9.1, HOT2) and were brought to a final volume of 80 μl with nuclease free water in a 200 μl thin wall PCR tube. We took 1 μl of each experiment to perform a quality control using Agilent 2100 bioanalyzer with the RNA 6000 Pico LabChip® Kit (Agilent) and NanoDrop 1000 Spectrophotometer (Thermo Scientific). Once we confirmed that RNA samples were not degraded, they were denatured at 90°C for 2 min (in the thermal cycler with heated lid on) and the tubes were immediately placed on ice for 2 min. Then, 10 μl of ice-cold 10X RNA structure buffer (Ambion) were added to samples and mixed by pipetting up and down several times. RNA samples were subsequently brought from 4°C to 23°C, in 20 min (1°C per min) in a thermal cycler. Finally, 10 μl of nuclease free water, and corresponding dilutions of RNase V1 (Ambion) or S1 nuclease (Fermentas) were added to the samples inside the thermal cycler for non-digested, V1-digested and S1-digested samples, respectively (see the following section “Determination of the optimal RNase V1 and S1 nuclease concentrations”). After mixing by pipetting, samples were incubated at 23°C for 15 minutes. The RNAs were purified using RNeasy MiniElute Cleanup kit following manufacturer’s instructions (Qiagen). We took 1 μl of each experiment to perform a quality control as described earlier (Supplementary figure S2B-D).

For the probing with RNase A enzyme, we included in a total of 2 μg of PolyA RNA from *S.cerevisiae* 20 ng of the following RNA molecules: TETp4p6, TETp9-9.1, SRA, B1 and U1. All the experiment was performed following exactly the same steps previously described, but adding 0.05 μg of RNase A (Ambion) in the samples instead of V1 or S1 enzymes.

#### Determination of the optimal RNase V1 and S1 nuclease concentrations

The original PARS protocol used 0.01 U of RNase V1 (Ambion) and 1000 U of S1 nuclease (Fermentas) in a 100 μl reaction volume in their first study (Kertesz et al. 2010), which the authors claimed are the appropriate enzyme concentrations for cleavage reactions occurring with single-hit kinetics. However, in their next study (Wan et al. 2013) they suggested a titration of nucleases to choose the optimal conditions for cleaving RNA molecules once per molecule on average, to avoid putative conformational changes after the first enzymatic cleavage. In their work they considered that this single-hit kinetics happened when around 10-20% of the RNA is cleaved. The authors suggested that the titration of nucleases could be done radiolabeling the RNA molecules, digesting them with different enzymatic dilutions, running them on a gel and quantifying the percentage of full-length RNA before and after the digestion. In our study, we went for a more direct approach performing full nextPARS experiments using different enzyme dilutions and probing some RNA control molecules with known secondary structure. In this wasy, we tested different dilutions of both enzymes and checked their bioanalyzer profile with the RNA 6000 Pico LabChip® Kit (Agilent). This served to assess the digestion pattern and confirm that RNA samples were not completely digested or not digested at all (Supplementary figure S2C,D). Then, rather than measuring the percentage of undigested/digested RNA molecules as in previous studies developing PARS technique (Wan et al. 2013), we directly analyzed the sequencing results of different samples treated with distinct enzyme concentrations, as well as a non-digested sample. In this way, we could directly assess the optimal enzyme concentration that resulted in a digestion profile that gives more accurate results according to the known secondary structure of positive controls. Specifically, we tested the following RNase V1 dilutions (number between parentheses correspond to units used in a 100 μl reaction volume): 1:30 (0.03 U), 1:50 (0.02 U), 1:100 (0.01 U) and 1:250 (0.004 U). We also tested 1:10 (0.1 U) and 1:20 (0.05 U) RNase V1 dilutions, but samples were completely digested, so we did not proceed further to the preparation of the libraries. For S1, the following dilutions were tested: stock concentration (1000 U), 1:2 (500 U), 1:5 (200 U), 1:20 (50 U), 1:50 (20 U). Triplicates were performed for all samples. The final concentrations used for our experiments were 0.03 U and 200 U for V1 and S1, respectively.

### Library preparation

#### nextPARS: Library preparation using TruSeq Small RNA Sample Preparation Kit (Illumina)

With the aim to have higher sequencing throughput and multiplexing capacity, we implemented an adapted PARS protocol to Illumina sequencing platform to study genome-wide the secondary structure of RNA molecules, which we named “nextPARS” (Figure1). In PARS, after the *in vitro* folding and RNase digestion step, the authors include a random fragmentation step and a size selection of RNA fragments by a column cleanup that we skip in the nextPARS protocol. In PARS, the first ligation is the 5’adapter and the second one the 3’adaptor after a phosphatase treatment. In nextPARS, we performed a phosphatase and a kinase treatment just after the enzyme digestion, to leave a hydroxyl group at the 3’ end and a phosphate group at the 5’ end of all RNA fragments coming from nuclease digestion to ligate them to the adaptors in the further library preparation steps. To control for unspecific degradation of RNA, we included the non-digested sample. For the phosphatase treatment, we incubated at 37°C for 30min a reaction mix with 16 μl of the non-digested, V1- and S1-digested samples, 2.5 μl of 10X phosphatase buffer, 2.5 μl of nuclease-free water, 1 μl of RNAse inhibitor and 3 μl of Antarctic phosphatase (New England BioLabs Inc.). After 5 minutes at 65°C, we put samples on ice and added 4 μl of T4 Polynucleotide Kinase (PNK, New England BioLabs Inc.), 5 μl of 10X PNK buffer, 10 μl of ATP 10 mM, 1 μl of RNAse inhibitor and nuclease-free water up to a total volume of 50 μl. After 1 hour of incubation at 37°C, samples were purified using RNeasy MiniElute Cleanup kit following manufacturer’s instructions (Qiagen) with a 10 μl RNase-free water final elution step. Then, samples were concentrated using a centrifugal evaporator Speed Vac® to a final volume of 5 μl and we started the TruSeq Small RNA Sample Preparation Kit (Illumina) protocol. All reagents used in the next step are from the Illumina kit if not specified otherwise.

Briefly, we first performed the 3’ adapter ligation with an initial denaturing step at 70°C for 2 min with the 5 μl of RNA samples and 1 μl of RNA 3’ adapter. Samples were then put on ice, and 2 μl of 5X HM Ligation Buffer, 1 μl of RNAse inhibitor and 1 μl of T4 RNA Ligase 2, truncated (New England BioLabs Inc.) were added. After 1 hour incubation at 28°C we added 1 μl of stop solution, gently pipetted up and down, incubated for 15 min more at 28°C and placed the samples on ice. Next, for the 5’ adapter ligation, we denatured the RNA 5’ adapter (1.1 μl per sample) at 70°C for 2 min and placed the tube immediately on ice. We added 1.1 μl of 10 mM ATP and 1.1 μl of T4 RNA Ligase per each sample in the same tube, mixed it, and added 3 μl of the mix in each of the samples coming from the 3’ adapter ligation. We incubated them at 28°C for 1 hour. We then performed the reverse transcription of the samples starting with a denaturing step at 70°C for 2 min of 6 μl of the 5’ and 3’ adapter-ligated RNA with 1 RNA RT primer. After putting the samples on ice, we added 2 μl of 5X First strand buffer, 1 μl of SuperScript II Reverse Transcriptase (Life Technologies), 1 μl of 100 mM DTT, 1 μl of RNAse inhibitor, and 0.5 μl of 1:2 diluted dNTP mix and incubated them at 50°C for 1 hour. To perform the PCR amplification we added to the samples 8.5 μl of ultra-pure water, 25 μl of PCR mix, 2 μl of RNA PCR primer and 2 μl of RNA PCR primer indexed (with a different index in each of the samples tested). Cycling conditions began with a denaturation step of 30 seconds at 98°C, followed by 11 cycles of 10 seconds at 98°C, 30 seconds at 60°C, and 15 seconds at 72°C, with a final extension step at 72°C for 10 min. We diluted 1 μl of each sample in 1 μl of ultrapure water to perform a quality control using Agilent 2100 bioanalyzer with the High Sensitivity DNA Kit (Agilent).

Finally, we purified and size-selected the prepared libraries to get rid of primers and adapters dimers using Novex TBE 6% gel (Invitrogen). We loaded into the gel the total volume of cDNA constructs (50 μl), as well as the High resolution ladder and Custom ladder (1 μl each), mixed with the proper amount of DNA loading dye, and ran it at 145 V for 1 hour. Gels were stained for 10 min with 4 μl SYBR Safe (Invitrogen) mixed with 50 ml of TBE. Using blue light (Dark Reader; Clare Chemical Research), we cut a gel slice from around 180 to 500 nt, which was shredded as previously mentioned, and 400 μl of ultrapure water were added to elute the DNA by rotating the tube at room temperature for at least 2 hours. Both the eluate and gel debris were transferred to a Spin-X centrifuge tube filter (pore size 0.45 μm, Sigma-Aldrich) and centrifuged at 8000 rpm for 1 min. Then, 40 μl of 3M NaOAc, 2.5 μl of glycogen and 1300 μl of pre-chilled 100% ethanol were added and centrifuged at maximum speed for 20 min at 4°C. We washed the pellet with 500 μl of pre-chilled 70% ethanol and after centrifuging at maximum speed for 2 min and removing the ethanol, the pellet was dried placing the tube with the lid open in a 37°C heat block for around 10 minutes. We resuspended the pellet in 10 μl of EB buffer for at least 10 minutes and a final quality control of each library was run using Agilent 2100 bioanalyzer with the DNA 1000 Kit (Agilent).

### Sequencing

Libraries were sequenced in single-reads with read lengths of 50 nucleotides in Illumina HiSeq2000 machines at the Genomics Unit of the CRG. All raw sequences used in this project has been deposited in Short Read Archive (SRA) database under the project number PRJNA380612.

### Mapping of Illumina reads and determination of enzymatic cleavage points

Illumina reads were aligned with tophat2 version 2.0.9 with the --no-coverage-search option enabled (Trapnell et al. 2009). SOLiD reads from the original PARS publication (Kertesz et al. 2010) were aligned with SHRiMP version 2.2.3 (Rumble et al. 2009). We used several reference sequence sets depending on data analysed: *S. cerevisiae* S288C full chromosomes were downloaded from the *Saccharomyces* Genome Database (SGD) (Cherry et al. 2012), and we concatenated the sequences of the corresponding control molecules spiked to each experiments. Reads aligned non-uniquely (i.e. having mapping quality below 20) were ignored. Subsequently, enzyme cleavage positions were determined as follows. For each read alignment, we retrieved the 5’-end position in the reference genome, and compared this to the genome annotation. If the position coincided with exonic regions of the genome, the information about the cleavage site was stored. The resulting digestion profile is stored as the number of cuts per position of the transcript. The load is defined as the average number of cuts per position. We provide all necessary scripts to perform this operation in the following repository (https://github.com/Gabaldonlab/nextPARS).

### Assessment of correlations between digestion profiles

We compared all sequencing runs in a pairwise manner in terms of number of enzymatic cleavage points (cuts) for all transcript positions. For each pair of sequencing runs, we retrieved the number of cuts in all positions from all transcripts meeting the threshold described below and computed the Spearman’s correlation coefficients to ensure that the results were consistent. Since each run of the experiment will produce different values, and transcript expression levels may differ each time, we used the rank-based Spearman’s value to see if coinciding positions have the same relative number of cuts for the given enzyme. We defined the transcript load as the average number of inferred enzymatic cuts per position in a given transcript. We expect the load to depend on the relative concentration of the transcript in the sample and the depth of the sequencing run. Correlations at different load cut-offs are shown in Supplementary table S3. From this, we determined that a load of 5.0 or greater is optimal to retain a sufficient number of transcripts that all have a high enough expression level and sequencing depth to produce consistent results, so for subsequent analyses, transcripts with a load below 5.0 (on average less than five cuts per position of a transcript) in a given run were ignored. We computed correlations among replicates within each nextPARS and PARS experiment, as well as between the two approaches. Since nextPARS uses 50 nucleotide reads, the final 50 of each probed molecule are uninformative and are not included in the correlation calculations involving nextPARS (correlations for the original PARS with itself do include the final 50 positions). Table 2 shows the average correlations for the whole set of yeast transcripts as well as for the three control molecules shared by the different experiments (TETp4p6, TETp9p9.1, HOT2), while Supplementary table S1 shows all correlations results for the full dataset.

### Computation of nextPARS scores

In brief, nextPARS scores are derived as follows:

i. The input is a digestion profile which indicates the raw number of enzymatic cuts per position. One such profile is available for each enzyme and replicate.
ii. The raw numbers are capped to a given maximum percentile (the default is 95%, meaning the upper 5% of the values are set to the value at the 95th percentile). This step is introduced because there are often positions that are preferentially cleaved with numbers of cuts that are orders of magnitude greater than other positions.
iii. Then read counts from each digestion in a given molecule are normalized to its average, giving an average of 1 read per position in each molecule and run of the experiment. This is to account for a number of factors, including the different expression levels of each molecule, the potentially different sequencing depth between each run, and the different cutting rates between the S1 and V1 enzymes, allowing for a comparable range of values amongst all of the digestions. When comparing the read counts per site between S1 and V1 experiments, including only those molecules shared between both PARS and nextPARS datasets having an average of at least 5 counts per site (420 molecules in total), a Student’s t-test showed that in the PARS experiments, S1 experiments had significantly greater counts per site than V1 experiments (p=5.1e-14), while the reverse was true for nextPars (p=2.2e-16). So it is important to have a normalized value before comparing V1 against S1. Also, since part of the nextPARS protocol involves mapping reads with a length of 50 nucleotides, the final 50 sites have no information, and so these sites are given scores of 0 and not considered when normalizing to the average.
iv. When replicates are available, we obtain a single list of values for both V1 and S1 digestions, by taking the average at each position for all of the normalized V1’s and S1’s, respectively.
v. With these two lists we can calculate combined scores per position in a manner similar to the original PARS protocol, but now with a more reliable footing. Since the aim is to use these scores to determine whether each position is paired or unpaired, we must compare to a given threshold, and thus we will need a standard range of values for the scores. After the steps taken so far, there are no set maximum or minimum values for each position in either the V1 or S1 lists, so the combined scores must be normalized. A few different methods have been tested: A) subtract the S1 from V1, then normalize to give a maximum value of 1.0. B) Normalize V1 and S1 values, separately, to a maximum value of 1.0, then subtract S1 values from the corresponding V1 values, so that positive values would suggest a paired site and negative values would suggest an unpaired site. However, since we want to be able to apply a universal threshold value to scores when determining structural constraints, we need to have a fixed range of values (ideally from -1.0 to +1.0). Method B will generally have bounds smaller than this range because at most sites there are cuts from both enzymes. So we then tried method C), which is to first subtract S1 from V1 values, then normalize positive values to have a maximum of 1.0 and negative values to have a minimum of -1.0. With this method, there is still the potential bias toward one enzyme or the other (as S1 often has higher values than V1). So we tested method D), which essentially combines B and C by first normalizing S1 and V1 values to a maximum of 1.0, subtracting, and then normalizing positive values to 1.0 and negative values to -1.0, to ensure that sites with the strongest V1 score to always have a score of 1.0 and those with the strongest S1 scores to always have -1.0. The effect is not large, since the sites cut most frequently by one enzyme are typically cut infrequently by the other, but it ensures the appropriate range for the final scores.

The justifications for the normalization technique and the maximum percentile cap are shown in Supplementary figure S3. The final normalization method chosen was D. After normalization we produce a single nextPARS score. By applying a threshold, this score can be converted into a structure preference profile (SPP), which is modeled from that used in the SeqFold protocol (Ouyang et al. 2013). For each position in the molecule, if the combined score is greater than the threshold, it is assigned a value of 0, which indicates a paired site. If the score is less than the negative of the threshold, it is assigned a value of 1, for an unpaired site. Otherwise it is assigned “NA” as there is not enough information at that site to say definitively whether it should be paired or unpaired. With the maximum score being +/- 1.0, we use a default score threshold of 0.8 to obtain the optimal constraints. The full computational pipeline to derive nextPARS scores and structural constraints from raw nextPARS data can be found at (https://github.com/Gabal-donlab/nextPARS).

#### Accuracy on constrained structures

We used 5 control structures (TETp4p6, RDN5-1, RDN18-1, RDN25-1 and RDN58-1). The PDB IDs for their structures are noted above in the results section. The structures provided by PDB contain 3D coordinates of the molecules, so we converted these to connectivity table files (which represent secondary structures) using the RNApdbee webserver (Antczak et al. 2014). For these structures we compared the PARS-constrained structures, as well as nextPARS-constrained, to the reference structures from PDB, using RNAfold (Lorenz et al. 2011) with thresholds for our scores of >= 0.8 for double-stranded (ds) and <= -0.8 for single-stranded (ss). PARS-constrained structures were acquired from the supplementary material of the original PARS paper (http://genie.weizmann.ac.il/pubs/PARS10/pars10_catalogs.html).

## ACKNOWLEDGEMENTS

We thank Howard Chang’s lab (Stanford University School of Medicine) for providing us with the *Tetrahymena* ribozyme and the HOTAIR clones described in this paper. We acknowledge the support of the CERCA Programme / Generalitat de Catalunya. Finally, we thank Jose Luis Rodriguez-Alés and Damian Loska, and the Genomics Unit of the CRG (Centre for Genomic Regulation) for their technical assistance.

This work was supported by the Spanish Ministry of Economy and Competitiveness grants, ‘Centro de Excelencia Severo Ochoa 2013-2017’ [SEV-2012-0208]; the European Regional Development Fund (ERDF) from the European Union and ERC Seventh Framework Programme (FP7/2007-2013) [ERC-2012-StG-310325]; and the European Union’s Horizon 2020 research and innovation programme under the Marie Sklodowska-Curie grant agreement [H2020-MSCA-ITN-2014-642095].

